# The genomic footprint of social stratification in admixing American populations

**DOI:** 10.1101/2022.11.16.516754

**Authors:** Alex Mas-Sandoval, Sara Mathieson, Matteo Fumagalli

## Abstract

Cultural and socioeconomic differences stratify human societies and shape their genetic structure beyond the sole effect of geography. Despite mating being limited by the permeability of sociocultural stratification, most demographic models in population genetics often assume random mating. Taking advantage of the correlation between sociocultural stratification and the proportion of genetic ancestry in admixed populations, we sought to infer the former process in the Americas. To this aim, we define a mating model where the individual proportions of the genome inherited from Native American, European and sub-Saharan African ancestral populations constrain the mating probabilities through ancestry-related assortative mating and sex bias parameters. We simulate a wide range of admixture scenarios under this model. Then, we train a deep neural network and retrieve good performance in predicting mating parameters from genomic data. Our results show how population stratification shaped by racial and gender hierarchies have constrained the admixture processes in the Americas since the European colonisation and the subsequent Atlantic slave trade.

## Introduction

Assortative mating is the phenomenon whereby mates resemble each other more than would occur under random mating. Through the lens of population genetics, assortative mating entails the correlation of genetic variants between mates ***Versluys et al. (2021*)**. Assortative mating is often a consequence of a subdivided population structure, observed in most human populations ***Nagoshi et al. (1990); Sebro and Risch (2012); Sebro et al. (2017*)**. In many species, assortative mating is also the product of active mate choice, an adaptive behaviour by which individuals choose a genetically similar mate by their phenotype ***Merrill et al. (2019); Versluys et al. (2021*)**. Whether or not active mate choice significantly takes place in human populations is unclear, as it is challenging to discern the effect it might have beyond that of population structure ***Xie et al. (2015); Sebro and Risch (2012); Abdellaoui et al. (2014); Sebro et al. (2017*)**.

Geography structures populations of most species, including humans. Individuals separated by shorter distances interact more than individuals separated by longer distances or geographical features like watercourses or mountain ranges, which imply limited mating as the geographical barriers increase ***Wright (1943); Malecot (1948); Kimura and Weiss (1964); Cavalli-Sforza et al. (1996); Novembre et al. (2008*)**. The reduced gene-flow between geographical groups leads to differentiated processes of genetic drift and results in distinguishable genetic profiles known as genetic ancestries ***Mathieson and Scally (2020); Lewis et al. (2022); Coop (2022*)**.

In human populations, culture also shapes the population structure. Socioeconomic and cultural barriers, which might have a certain permeability, limit the interaction between human groups. These social groups with a common language, religion, socioeconomic status, etc define overlapping subpopulations where mating takes place mostly within them. ***Manni (2010); Campbell (2015); Matsumae et al. (2021*)**.

In the case of migration, two or more populations cohabit in the same location and eventually admix to become subpopulations of a newly admixed population. The admixture process does not take place randomly but it is limited by barriers set by socioeconomic and cultural differences between subpopulations with their own distinguishable profile of genetic ancestry. ***Risch et al. (2009); Nagoshi et al. (1990); Sebro et al. (2017*)**. These barriers are socially constructed and, particularly in colonial contexts, their permeability is often politically restricted ***McLean (2021*)**.

In admixed individuals, the genetic ancestry related to each source population can be tracked along the genome and expressed in individual-based proportions. Therefore, in recently admixed populations, the population structure driven by culture and socioeconomic differences is associ-ated with differences in the proportions of genetic ancestry. As a consequence, the proportion of genetic ancestry between mates correlate. This phenomenon is defined as ancestry-related assor-tative mating ***Burrell and Disotell (2009); Bryc et al. (2010); Norris et al. (2019*)**. In addition, when mating is not random regarding the ancestry proportions, individuals might show a preference towards mating partners of the opposite sex with lower or higher ancestry proportions, which is defined as ancestry-related sex bias ***Goldberg and Rosenberg (2015*)**.

In admixed populations, the length of the continuous ancestry tracts is widely used to infer the time since admixture under the assumption of random mating. During gametogenesis in individ-uals admixed individuals, recombination breaks down continuous ancestry tracts inherited from each of the source populations of the admixture event into smaller alternate fragments at each generation. Consequently, the length of the continuous ancestry tracts reflects how many generations ago the source populations migrated across the geographical barriers that prevented them to mate ***Gravel (2012); Hellenthal et al. (2014); Chintalapati et al. (2022*)**. Herein, we postulate that the tract length information can also monitor the permeability of socioeconomic and cultural barriers between subpopulations with different genetic ancestries.

Some of the methods to date admixture can discern multiple pulses of migration. Only a few of them have addressed complex admixture histories such as the fluctuation of unbalanced migra-tions of males and females from two source populations ***Laurent et al. (2022*)**. However, almost all these approaches assume random mating in the admixed population, overlooking the effect of population stratification in the population structure. To our knowledge, few studies have modelled ancestry-related assortative mating during admixture, although limited to two source populations ***Goldberg et al. (2020); Kim et al. (2021*)**.

Beyond analytical modelling, population genetics studies have also measured ancestry-related assortative mating through the correlation of genetic ancestry proportions between mates ***Bryc et al. (2010); Korunes et al. (2022); Arauna et al. (2022*)**. Non-random mating can also be monitored through deviations of the observed heterozygosity from Hardy-Weinberg equilibrium expected values ***Crow and Felsenstein (1968*)**. Thus, when information on genetic ancestry of mating couples is not available, it is still possible to infer ancestry-related assortative mating through the comparison of the genetic ancestry of the two homologous chromosomes of the individuals ***Norris et al. (2019*)**. Nevertheless, despite these efforts, we still lack a rigorous and robust method to shed light onto the patterns of ancestry-related non-random mating. More specifically, we are in need of a comprehensive model of ancestry-related assortative mating and sex bias, two parameters which have been rarely jointly modelled in population genetics.

Among human populations, the admixing populations from the Americas are of special interest in admixture studies. We consider them as *admixing* populations, because their genetics is shaped by an ongoing admixture process of three differentiated continental ancestries that started five centuries ago, constrained by a strong social structure.

At the end of the 15*^th^* century, European powers initiated the colonisation process in the lands inhabited by Native Americans. In this frame, European colonisers enslaved more than 10 million people brought from sub-Saharan Africa ***Eltis (2018*)**. As a result of this historical event, the populations of the Americas are the outcome of the admixture of Native American, European and sub-Saharan genetic ancestries ***Salzano and Bortolini (2005); Bedoya et al. (2006); Wang et al. (2008); Moreno-Estrada et al. (2013); Gravel et al. (2013); Ruiz-Linares et al. (2014); Adhikari et al. (2016, 2017*)**.

After the abolition of slavery, most of these populations kept stratified based on the socioeconomic status and according to hierarchical notions of racial difference. Some of them have even experienced explicit segregation policies long after the abolition that aimed to prevent mating between subpopulations from different origins and maintain socioeconomic stratification ***Douglass (1882); Du Bois et al. (1935); Davis (1981*)**.

In Latin America, in addition to segregation, European colonial powers and creole elites implemented eugenicist policies under the frame of *mestizaje/mestiçagem. Mestizaje/mestiçagem* refers to the process of admixture of Native American, European and sub-Saharan ancestries in the context of the European colonisation and therefore associated to the mixture across hierarchical differences understood as “racial”, differences of class and differences of gender. Since mid-nineteenth century, Latin American nation-building elites have aimed to associate *mestizaje/mestiçagem* to an equalising process, by claiming that it overcomes and blurs the socioeconomic differences related to “race”. However, critics have argued that the *mestizaje/mestiçagem’s* notion of hybridity inherently entails the idea of its constitutive origins and the hierarchies that order those origins. In this sense, *mestizaje/mestiçagem* attaches greater value to the interactions that move towards whiteness and masculinity and lower value to those that move towards blackness or indigeneity, and femininity. ***Wade (2017, 2020); Abel (2021*)**.

By analysing the impact of the European colonisation in the population structure through mating, we aim to evaluate the stratification related to the genetic ancestry not only quantifying the population subdivision but also measuring the genetic ancestry asymmetry between males and females in mating. Following this approach, we conceptualise a novel mechanistic mating model that explicitly integrates ancestry-related assortative mating and sex bias jointly, through an inter-sectional approach derived from the interrelated hierarchies observed in the admixture process. ***Crenshaw (1989, 1991); Wade (2017*)**. We consider a three-way admixture scenario mirroring the demography of the admixing American populations. We build and train a deep neural network to infer non-random mating parameters using extensive synthetic data. We deploy this network to genomic data from admixing American populations sequenced as part of the 1000 Genomes Project ***Consortium et al. (2015)*** and quantify the extent of ancestry-related assortative mating and sex bias. Finally, we discuss racial and gender hierarchies as inferred from their footprint on genetic structure.

## Results and discussion

We report our results in three sections: (i) the novel mating model and framework for simulations, (ii) the performance of the neural network, and (iii) the inference of the ancestry-related mating probabilities for admixing American populations.

### An ancestry-related mating model

We present an admixture model defined by the mating probabilities of all possible male and female couples, set by their ancestry proportion difference. For each ancestry, the ancestry-related sex bias (***SB***) and the ancestry-related assortative mating (***AM***) parameters determine the mating probability of each couple as a function of the difference in the ancestry proportion between male and female. We assume that the differences in the ancestry proportions within the mating couples follow a Normal distribution that translates into the mating probabilities (***Figure 1***).

**Figure 1.**
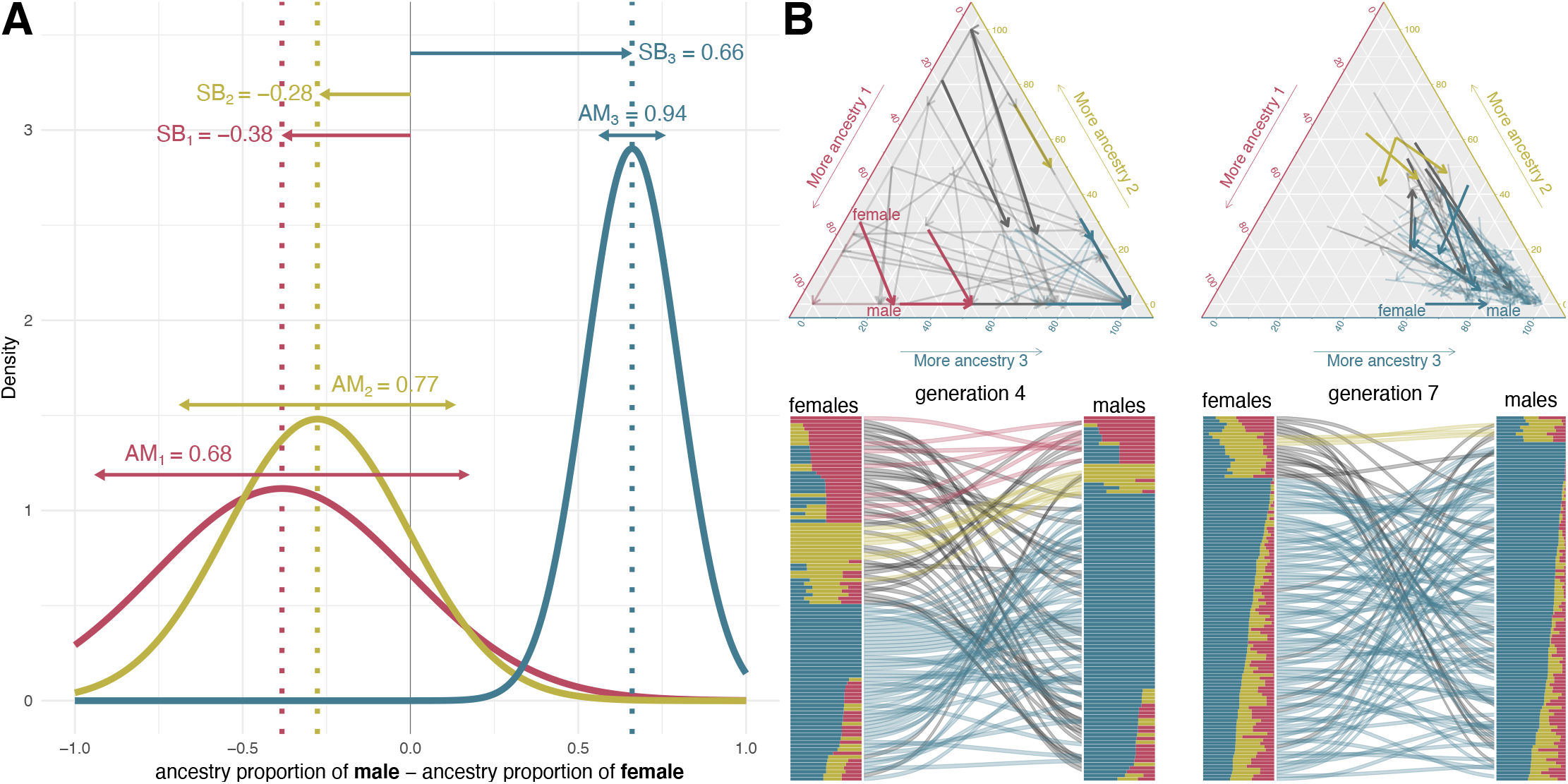
**A**: ***AM*** and ***SB*** values that modulate the mating probabilities in a simulation example of 19 generations from the colonization of America to nowadays. The mating probability for a given couple is set as a function of the differences in the genetic ancestry proportions for each ancestry. We assume the mating probability follow a three dimensional normal distribution. In this normal distribution, ***SB*** sets the expected value and ***AM*** is inversely proportional to its variance. **B**:Ancestry proportions of mating couples at generations 4 and 7 in ternary plots (top) and barplots (bottom) based on the mating probabilities defined in A.. In the top plots, each arrow represents a couple. The arrow tail and head coordinates in the ternary plots show the ancestry proportions of the female and the male, respectively. In the bottom, the barplots represent male and female ancestry proportions, linked by curved lines reflecting mating. Red, yellow and blue correspond to ancestries 1, 2 and 3. The arrows in the ternary plot and the lines between barplots representing a mating couple are coloured with the colour corresponding to the predominant ancestry in both male and female, and they are depicted in black if it differs within between them.

The expected value of this Normal distribution defines ***SB***, while ***AM*** is modelled as being in-versely proportional to the variance (see Methods and Materials for mathematical details). For a given ancestry, positive ***SB*** values indicate that couples where males have a higher proportion than females of this ancestry have more chances of mating, with the opposite pattern for negative ***SB*** values. Conversely, ***AM*** modulates the decay of mating probability when the difference of ancestry proportion within the couple moves away from the expected value set by the ***SB*** parameter. Therefore, when ***SB*** is close to zero, a couple with substantial differences in their ancestry proportions has a higher probability of mating if ***AM*** is lower (***Figure 1**A*). ***Figure 1**B* shows how a sample of individuals at generations 4 and 7 mate based on the mating probabilities set by the example values of ***AM*** and ***SB*** defined in ***Figure 1**A*.

We focus on the case of three-way ancestry, a model that describes the admixture of the populations of the Americas and their triple genetic ancestry: Native American, European and sub-Saharan African. We define two alternative models, referred to as the One Pulse model and the Two Pulses model. The simpler One Pulse model assumes one migration event occurring 19 generations ago and includes five independent parameters: ***AM***_1_ ***AM***_2_, ***AM***_3_ and ***SB***_1_ and ***SB***_2_, for sub-Saharan African (1), Native American (2) and European (3) ancestries (***Figure 1***A). In the Two Pulses model, an additional parameter (the Gene Flow Rate at generation 10 -*GFR*_*gen*10_-) determines the fraction of the gene pool arriving in a second migration from each source population at generation 10.

Our goal is to predict ***AM*** and ***SB*** (and *GFR*_*gen*10_ for the Two Pulses model) for the admixing American populations sampled in the 1000 genomes project (African Caribbeans in Barbados, ACB; African Ancestry in South-West USA, ASW; Colombians in Medellín, CLM; Mexicans in Los Angeles, MXL; Peruvians in Lima, PEL; and Puerto Ricans in Puerto Rico, PUR). To do so, we aim to compare the continuous ancestry tract lengths profile obtained from a local ancestry analysis performed on this data to the tract lengths profile issued from simulated data for each population with known combinations of the mating parameters.

For 10,000 random combinations of ***AM*** and ***SB*** parameters (and ***GFR***_*sen*10_ for the Two Pulses model) for each population we simulate, forward-in-time, a range of admixture scenarios. The contribution of each genetic ancestry to the gene pool of the simulated admixed population is equivalent to the observed ancestry proportions after the local ancestry analysis on the real data. We simulate 22 autosomal chromosomes and the X chromosome for each individual at each generation, keeping track of the local genetic ancestry at each chromosomal region (***Figure 1**, **Figure 2**A*). This approach serves a dual purpose: (i) to simulate the mating as a function of the genome-wide ancestry proportions of all males and females, based on the mating probabilities set by the ***AM*** and ***SB*** parameters of the mating model (***Figure 1***); (ii) to generate the continuous ancestry tract lengths profile as an output after the last simulated generation, which counts the number of fragments in 22 length windows in a logarithmic scale (***Figure 2***B).

**Figure 2.**
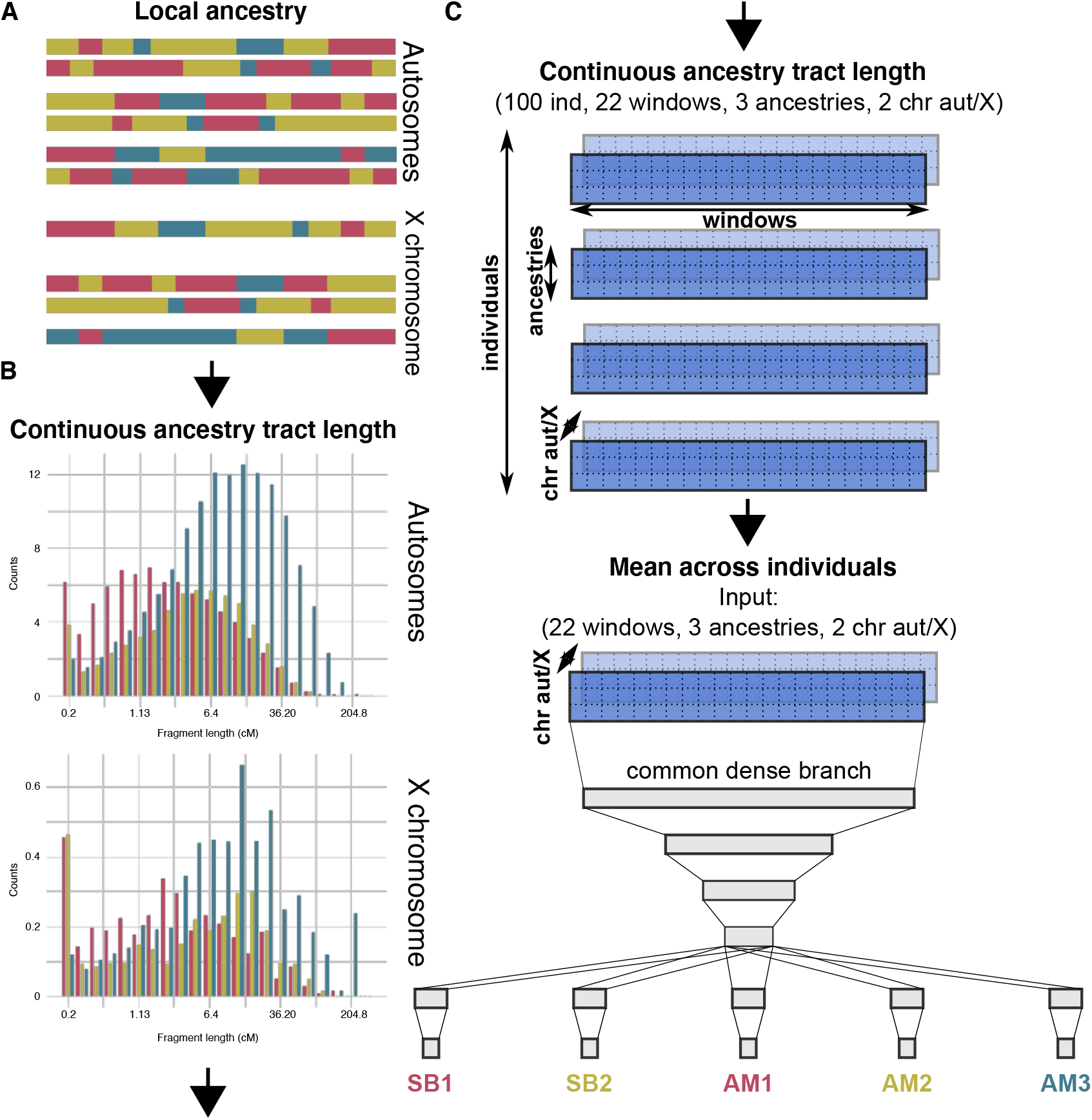
**A**:Schematic view of the autosomal and sex chromosomes split into the continuous ancestry tracts inherited from each of the three ancestries after a local ancestry analysis with RFMix. **B**:Continuous ancestry tract length profile displaying the number of tracts for each ancestry in each tract length bin. The break points that define the bin widths are set in a logarithmic scale. **C**: Matrix representing the continuous ancestry tract length profile accounting for the amount of tracts in each length bin, for each ancestry in either autosomal or sex chromosome for each indivdiual. The mean across individuals summarises the 4-dimensional matrix in a population 3-dimensional matrix, which is used as the input of the neural network. The neural network has four fully connected layers that split in a branch for each parameter, each one made of a last hidden layer connected to the output layer.

### A deep neural network to estimate mating parameters efficiently

To infer all parameters in our model, we train a deep neural network for each popualtion. By exploring the entire parameter space of ***AM*** and ***SB*** parameters (and ***GFR***_*sen*10_ for the Two Pulses model), we feed simulated continuous ancestry tract length profiles to a deep neural network consisting of fully-connected layers(Figure ***Figure 2***C).

The network sufficiently learn the weights for all parameters without overfitting over 40 epochs, as shown by the decay of the loss function (mean squared error) (***Figure 3-Figure Supplement 1***). We observe low mean squared error on the testing set for all parameters (***Figure 3***). Similarly, we appreciate a high correlation between true and predicted values, as shown by ***R***^2^ values and the confusion matrices, at testing (***Figure 3-Figure Supplement 2***).

**Figure 3.**
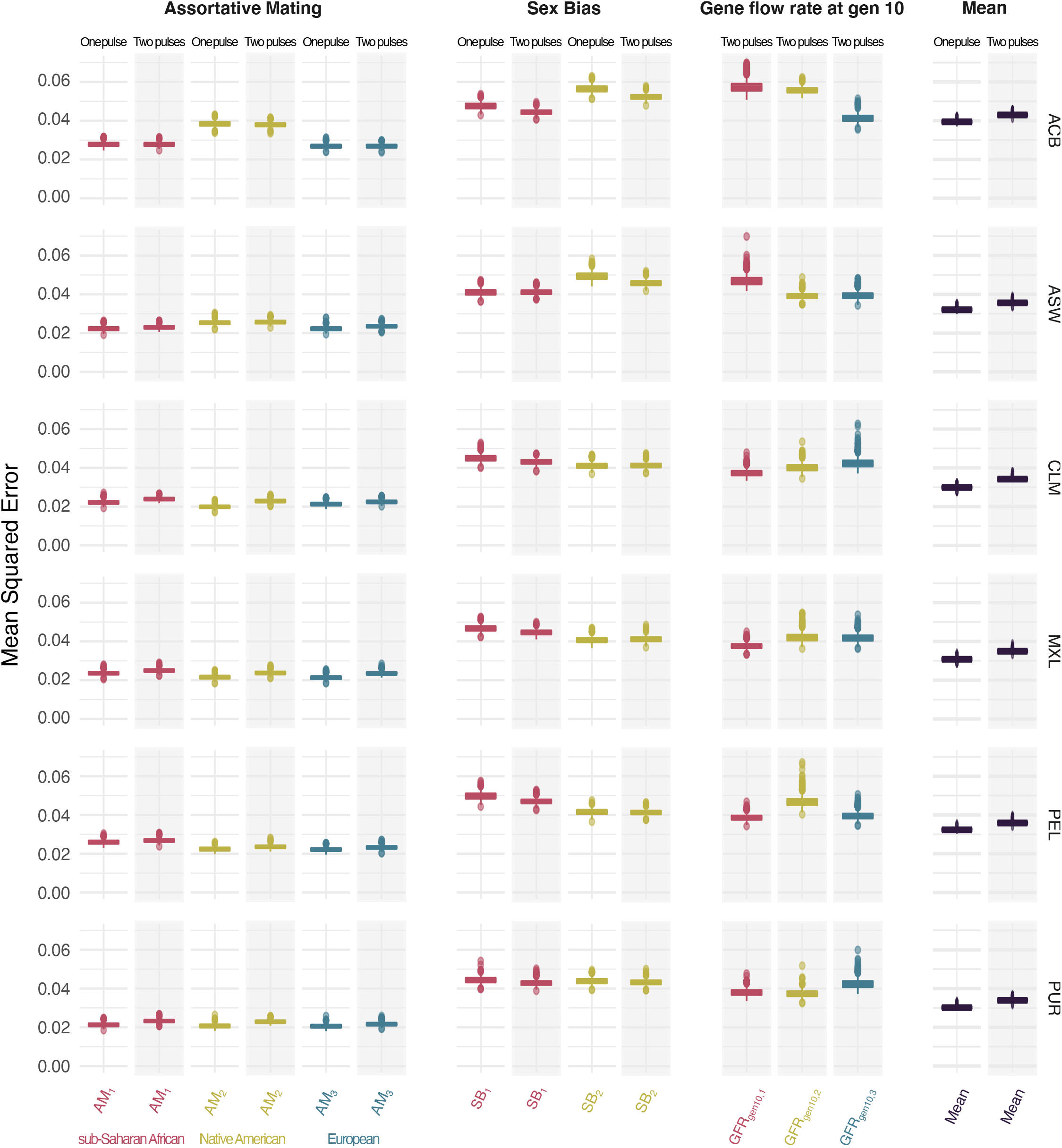
Mean squared error comparing true and predicted values at testing for the *Assortative Mating, Sex Bias, Gene flow rate at generation 10* parameters for each ancestry for both One Pulse and Two pulses models and mean values for each model. Each color represents the values for a different ancestry (red, yellow and blue for ancestries 1, 2 and 3 respectively, which correspond to sub-Saharan, Native American and European ancestries)

The trained network exhibits better predictions for ***AM*** parameters than for ***SB*** and ***GFR***_*gen*10_ parameters across all ancestries, populations, and migration models. Interestingly, the higher complexity of the Two Pulses migration model does not produce higher mean squared error or lower ***R***^2^ valuesfor any of the tested parameters. In fact, the mean of the mean squared error for the Two Pulses model is only marginally higher than the mean of the mean squared error for the simpler One Pulse model. (***Figure 3, Figure 3–Figure Supplement 1, Figure 3–Figure Supplement 2***).

### The Native American and sub-Saharan genetic ancestries respectively shape the mating probabilities in Latin American and African American populations

We sought to test the occurrence and extent of assortative mating and sex bias in the admixing American populations from 1000 genomes. To predict ***AM*** and ***SB*** parameters (and ***GFR***_*gen*10_ for the Two Pulses model) we deployed the trained neural network on the continuous ancestry tract length profiles these populations obtained after a local ancestry analysis.

In the One Pulse model, the Latin American populations (CLM, MXL, PEL, PUR) present a consistent pattern where the Native American ancestry shapes the mating probabilities, as the ***AM*** parameter associated to this ancestry is the highest in all populations. Thus, the differences in the Native American ancestry between males and females modulate the mating in Latin American populations, although the Native American ancestry is not the one observed in highest proportion in all of them. The high ***AM*** values are coupled with negative ***SB*** for CLM, PEL and PUR populations, indicating that females of high Native American ancestry are more likely to mate with males of lower Native American ancestry. MXL and CLM populations show stronger ***AM*** values than PEL and PUR, while PUR and CLM exhibit stronger sex-biased admixture than MXL and PEL. Conversely, ASW population presents the highest ***AM*** in the sub-Saharan African ancestry. Paired with a positive ***SB*** value, these estimates indicate that males of high sub-Saharan African ancestry are more likely to mate with females of lower sub-Saharan African ancestry. Finally, ACB populations show similar ***AM*** values for the three ancestries, with no specific ancestry modulating the mating probability (***Figure 4***A).

**Figure 4.**
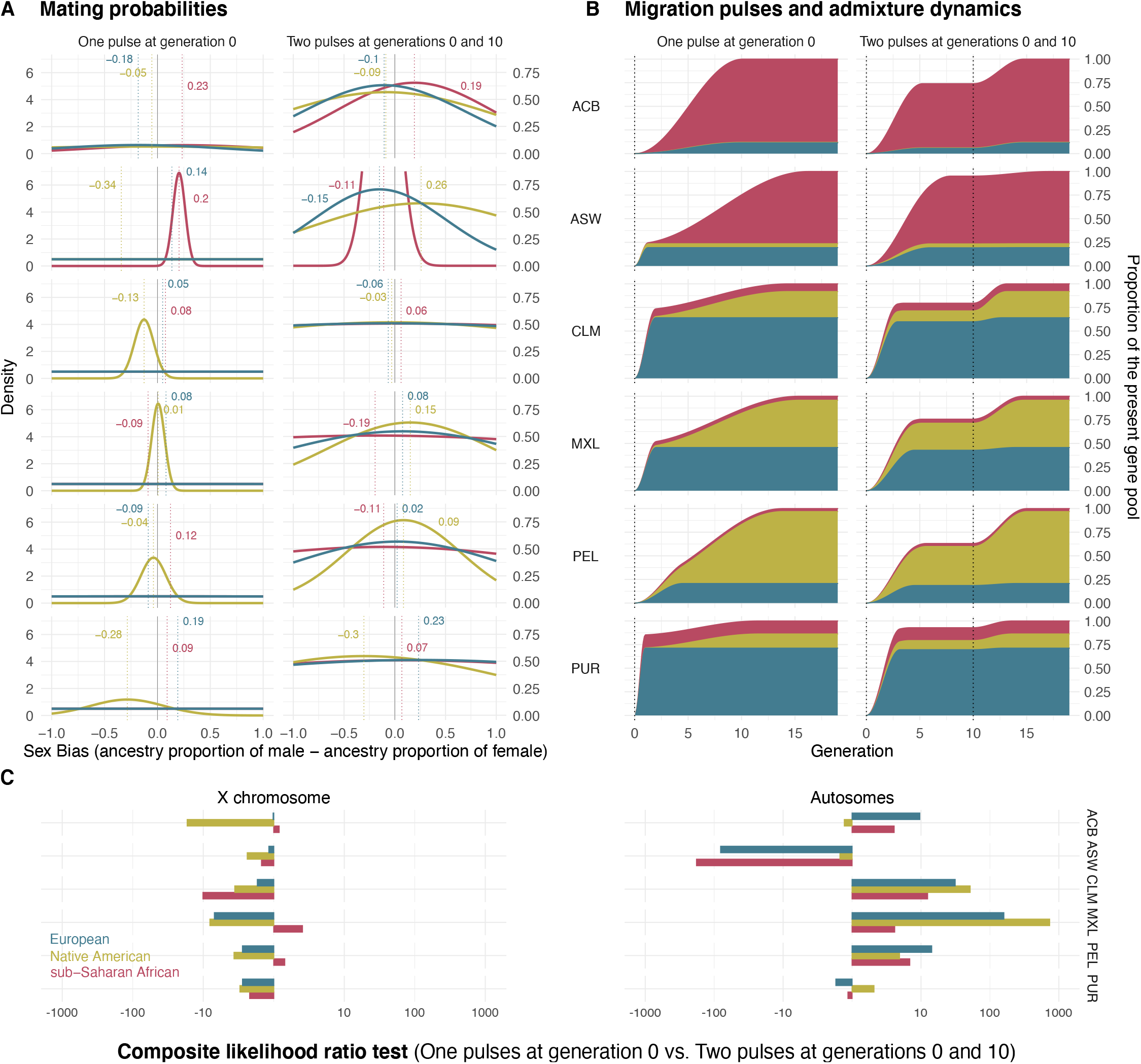
**A**: Mating probabilities as a function of male and female proportions of each ancestry, for each population. **B**: Migration pulses of each ancestry according to scenarios allowing one pulse and two pulses for each population. The y axis represents the cumulative increase in the ancestry specific gene pool relative to the final ancestry proportions, at each generation. The ancestry proportions at generation 19 represent the observed ancestry proportions of each population in real data. The increase in the cumulative ancestry specific gene pool is defined by ***GEN***_*gen*10_, while the slope of the increase is represented inversely proportional to ***AM***. **C**: Composite likelihood ratio test comparing two pulses model vs. one pulse model, for each ancestry for both the X chromosome and the autosomes. In this plot, positive values show a higher likelihood of the two pulses model based on the fit of the real fragment lengths in the distribution of fragment lengths of simulated data under the ***AM*** and ***SB*** parameters predicted by the neural network.

In the Two Pulses model, we allow for an additional migration pulse at generation 10 and we let the neural network predict the gene flow rate through the ***GFR***_*gen*10_ parameter. Under this new scenario, as expected, ***AM*** values are much lower than their corresponding values under a One Pulse model. In fact, part of the population structure that is modelled as social stratification in the One Pulse model is now modelled by gene flow from an additional migration event. Both models reflect similar admixture dynamics, where Native American and sub-Saharan African genetic components are gradually absorbed into Latin American and African American gene pools, respectively. (***Figure 1***B).

Under both models of migration, the effect that ***SB*** has on the mating probabilities depends on the ***AM*** values, as lower ***AM*** values imply a lower effect of ***SB*** on the mating. In case of low ***AM***, individuals are less constrained in their mating by their ancestry and, therefore, the effect of ***SB*** is less prominent.

We next sought to test whether observed genomic data is more compatible with a One Pulse or Two Pulses migration model. To this aim, we calculate the composite likelihood ratio to compare the fit of simulated continuous ancestry tract length profile under the predicted values of ***AM*** and ***SB*** (and ***GFR***_*gen*10_ for the Two Pulses model) to the empirical data. Results show that the Two Pulses model is more supported by the data for most populations in the autosomal chromosomes. On the other hand, the composite likelihood ratio suggests a better fit for the One Pulse model with stronger values for ***AM*** and ***SB*** in the X chromosome (***Figure 4**C*).

## Discussion

We tackle the analysis of the ancestry-related non-random mating driven by social structure through a mating model, which allows us to globally evaluate the forces that modulate population structure but also study the effect of these population dynamics at the individual level. Our results show evidence of ancestry-related sex bias and assortative mating in American admixed populations. In Latin Americans the proportion of Native American ancestry of men and women shape the mating probabilities and, therefore, the genetic structure of the population. By contrast, in African Americans, the sub-Saharan African ancestry modulates mating. Below, we discuss the significance of these results and the importance of our approach in studying social stratification. Finally, we evaluate the performance of our pipeline in discerning between migration and assortative mating and we explore how next steps could incorporate more complex admixture scenarios.

### Social stratification by racial and gender hierarchies

The ultimate aim of our approach is to infer social stratification in the Americas from the analysis of the population genetic structure. To this end, we model ancestry-related assortative mating mediating the extent of the effect of ancestry-related sex bias. We infer how they shape the mating dynamics of the population and we evaluate how these two dimensions affect at the individual level, thanks to the mating model framework.

We have defined this model from an intersectional perspective, which understands that racial, gender and class hierarchies are mutually constituted ***Davis (1981); Gonzalez (1983); Hooks (1984); Crenshaw (1989,1991); Collins (1990); Carneiro (1995); McCall (2005); Hancock (2007); Viveros Vigoya (2016)***. Besides, from Decolonial Feminism, authors have stressed the significance of the frame of the European Colonisation in redefining the concept of gender in America and have pointed to sexuality and mating as one of the contexts where racial, gender and class inequalities manifest with more visibility ***Stolcke (1992); Lugones (2007, 2008); Viveros Vigoya (2009***, 2016). Specifically, in our approach, we consider that racial stratification intensifies gender inequalities during mating. At the individual level, we conceive that the effects of these social hierarchies on mating depend on the relative position of the subject in racial and gender axes of inequality respect to the position of the other individuals of the population, and therefore on the *context.**Yuval-Davis (2006); Anthias (2012); Jorba and Rodó-Zárate (2019)***.

In this framework, we tackle the hypotheses about the role of *mestizaje/mestiçagem* in the inequality of a society developed by ***Wade (2017***, 2020) through a genetic study. Wade states that *mestizaje/mestiçagem* is a highly ambivalent discourse and set of practices, as it promotes and facilitates interactions across hierarchical differences of “race”, class and gender, but simultaneously reinforces those hierarchies. Studying the interaction of racial and gender hierarchies together, our approach allows us to investigate the dynamics of *mestizaje/mestiçagem* and challenge the oversimplified understanding that greater mixture translates into a decrease of discrimination and inequality. As such, high levels of mixture cannot be understood as a sign of equality or low discrimination if it takes place with strong gender biased patterns. Instead, this scenario provides evidence for a deep interaction between racial and gender hierarchies that shapes the social structure of the population.

In the specific case of the Latin American populations considered here, the prejudices and biases associated with being a woman of high Native American ancestry constrain the dynamics within the society and determine the mating, shaping the population genetic structure. The effect of these biases appears to be higher for racial hierarchies in MXL and CLM populations compared to PEL and PUR populations, while the effect of gender hierarchies appears to be stronger in PUR and CLM populations. The admixture dynamics, either modelled by assortative mating or migration pulses, show a slower and progressive absorption of the Native American ancestry into the admixed population. This is consistent with historical records reporting an increased pressure on Native populations to culturally assimilate towards a white/mestizo norm starting in the 19*^th^* century, which spurred internal migrations to urban areas and the loss of indigenous languages in places like Mexico and Peru ***Viqueira (2010); Telles (2014*)**.

Our results support the idea that *Mestizaje/mestiçagem* operates through racial and gender hierarchies and it is accompanied by a gradual dilution of the sociocultural elements associated with non-European genetic ancestries (found in higher proportions in mating females than in mating males) into a mainstream identity associated to the European gene pool (in which males contribute more than females).

To have a broader perspective of how racial and gender hierarchies operate across American societies, further approaches should analyse a wider dataset with more diverse, representative and carefully sampled populations. Specifically, the inclusion of socioeconomic variables in the sampling would allow us to evaluate how class hierarchies interrelate with racial and gender hi-erarchies. In addition, the transition to a more complex admixture model that monitors changes in the ***AM*** and ***SB*** parameters should provide the possibility to evaluate how racial and gender hierarchies have changed through history in different regions of the Americas.

### Discerning geographical barriers and social barriers

The disentanglement of social barriers from geographical barriers is a major challenge to face when addressing the complexity of admixture. Although non-random mating patterns associated with social structure and gene-flow after migration might leave similar footprints on the genome of a population, the distinction between these scenarios has important implications for understanding society.

The continuous ancestry tract length profile issued from our mating model is different from that issued from an admixture model where the admixed population only receives constant geneflow from migration of non-admixed source populations to model the population structure. In the latter approach, as defined in ***Goldberg and Rosenberg (2015); Laurent et al. (2022*)**, after migration the non-admixed source populations introduce full length non-recombined chromosomes with a single ancestry in the admixed populations and shape an identifiable pattern. In our approach, the continuous ancestry tract length profile is capturing this pattern to discern between ancestry-related assortative mating and gene-flow due to migration. Although this pattern characterised by low or non-recombined continuous ancestry tracts in a few individuals could remain hidden in a population mean continuous ancestry tract length profile, it should be detectable with and individual-based one.

Alternatively, a population structure correlated with genetic ancestry could also be modelled with an island model starting from non-admixed panmictic subpopulations with migration rates between them split by sex. Our approach resembles this model but considers a more realistic population continuously structured by genetic ancestry, rather than discrete (although permeable) subpopulation panmictic demes. In this sense, we address population structure through a mating model, which helps to focus the discussion on the effect at individual level of the dynamics that stratify the population.

A versatile mating model to accommodate a wider range of admixture scenarios The different patterns inferred in autosomal and X chromosomes suggest that some of the complexity of admixture is still not explained by our models. Whilst inferences based on autosomal chromosomes indicate a demographic scenario involving multiple migration events, inferences on the X chromosome support sex-biased admixture. Taken together, these trends suggest a need to expand the model by adding either the possibility of sex-biased migrations or changes in ***SB*** and ***AM*** through time. The flexibility of our methodology provides the basis to disentangle more complex scenarios of admixture in a further approach. The plasticity of forward-in-time simulators allows for additional complexity in the admixture model, by modifying the ***AM*** and ***SB*** parameters through time and modelling multiple migration pulses and population growth.

Machine learning has the potential to infer a large number of parameters issued from complex models of admixture. However, this step likely implies a shift from a population-based continuous ancestry tract length profile to an individual-based one, which needs to be linked to a redesign of the architecture of the neural network. Other architectures such as convolutional neural networks and generative models have been recently deployed to infer introgression, structure, admixture proportions, and post-admixture selection from population genomic data ***Gower et al. (2021); Wang et al. (2021); Meisner and Albrechtsen (2022); Hamid et al. (2022*)**.

In conclusion, an interdisciplinary approach that incorporates up-to-date insights from social sciences is essential to conceptualise population genetic models that aim to evaluate genetic structure driven by social stratification. Further studies might expand the analysis of population stratification by exploiting the full potential of machine learning in population genetics. An intersectional perspective that jointly addresses the effects of racial, gender and class hierarchies on population structure will be key to understanding the genetics of the admixing populations of the Americas.

## Methods and Materials

### A mating model defined by ancestry-related Assortative Mating and Sex Bias

We derived a mechanistic model where ***AM*** and ***SB*** parameters constrain the mating probabilities as a function of the difference in the ancestry proportion between male and female, for each ancestry. We assume that the probability of mating follows a normal distribution where ***SB*** for each ancestry is defined as the expected value of the difference in the ancestry proportion within mating couples, while ***AM*** is modelled as being inversely proportional to its variance (***Figure 1***).

#### Model definition

Consider an admixed finite population from ***S*** isolated source populations comprised of ***F*** females and ***M*** males. Assume that for each individual ***i*** we have a vector of inferred ancestry proportions 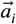 for each source population ***s***, so that 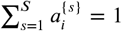. We consider a random variable for mating ***L*** as a realisation of the event *l_f,m_* between a female *f* and male ***m***.

We calculate the probability of mating between a female ***f*** and male ***m*** as

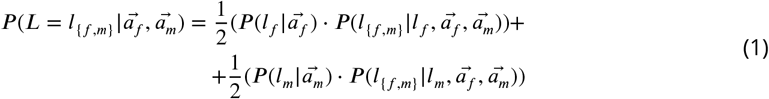

where 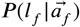 and 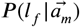 are the probabilities of either female or a male to start a mating event that will have one child as the outcome.

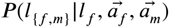 is the probability of a female to mate with a male given the ancestry proportions of both female and male, once the female has already been chosen to initiate the mating.

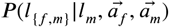is therefore the probability of a male to mate with a female given the ancestry proportions of both female and male, once the male has already been chosen to initiate the mating.

#### Mating probability of a couple

In the most basic model, all individuals can be assumed to have the same probability to start the mating event independent of their ancestry:

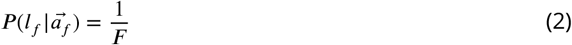

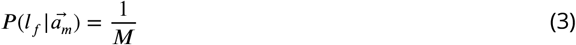

Once either a female or a male initiates the mating, the mating probability of each possible couple is defined as a function of the ancestry proportions of this individual and the ancestry proportions of each individual of the other sex 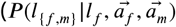 or 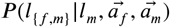 for respectively a male or a female that initiates the mating).

This probability is described by a multivariate normal distribution defined by a mean vector ***μ***, related to ***SB***, and covariance matrix Σ, related to ***AM***. This multivariate normal distribution consists of S variables defined as 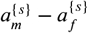, related to each ancestry *s*. They account for the difference of the ancestry proportion in each mating couple. The final mating probability for an individual and a possible mate is relative to the sum of all the probabilities for all the possible mates for this ndividual:

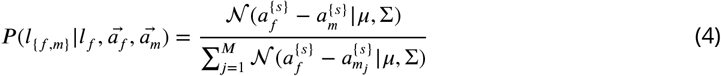

and:

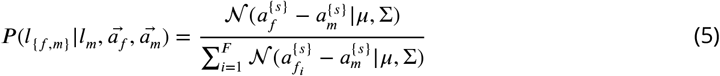

where ***μ*** is the vector of the expected means of the ancestry proportion differences (i.e 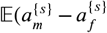) for ancestry ***s***) which define ***SB*** for each ancestry. The diagonal of Σ is the vector 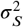, where the variance ***σ***^2^ of 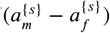 is inversely proportional to ***AM*** for each ancestry (***Figure 1***).

The sum of the mean vector (i.e. the sum ***SB*** parameters for all the ancestries) is zero 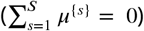. In addition, Σ is not full rank (|Σ| = 0). Consequently, the multivariate density function can be represented with ***S*** - 1 dimensions, which has 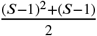 independent parameters.

#### The case of three ancestries. S=3

When S=3, the multivariate normal distribution is equivalentto a 2 dimensional multivariate normal distribution, which has five independent parameters (2 in ***μ*** and 3 in Σ):

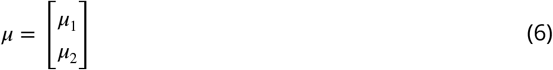

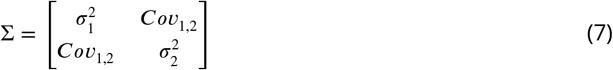

where ***Cov***_1,2_ can be defined by the variances of the three dimensional multivariate normal distri bution, including 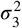:

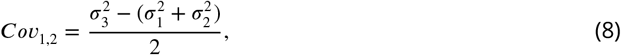

The mating model for three ancestries is set by sex bias for ancestries 1 and 2 (***SB***_1_, ***SB***_2_) and the assortative mating for ancestries 1, 2 and 3 (***AM***_1_, ***AM***_2_, ***AM***_2_). Therefore, ***SB*** for ancestries ***a*** ∈ {1,2} is defined as follows:

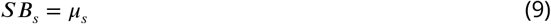

and ***AM*** for ancestries ***s*** ∈ {1,2,2}:

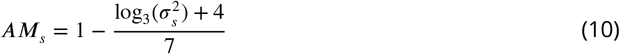

This arbitrary parameterisation has been chosen in order for the the assortative mating parameter ***AM_S_*** to cover the full spectrum of meaningful values taking values from 0 to 1 on a logarithmic scale, where 0 is random mating and 1 is very strong assortative mating.

### Simulations

We performed 10,000 simulations per population (CLM, MXL, PEL, PUR, ACB,ASW from 1000 genomes) following the mating model described above and exploring the joint parameter space of ***AM*** and ***SB*** (and ***GFR***_*gen*10_ for the Two Pulses model) using SLiM ***Haller and Messer (2019*)**. We simulated the 22 autosomal chromosomes and the X chromosome for each individual. We tracked their real local ancestry by recording the source population from which each genomic fragment is inherited. At each generation, we simulated mating based on the mating model as a function of the genetic ancestry proportions of the individuals. After the mating of two individuals, we simulated recombination in the gametogenesis of the offspring and the progressive break down of the continuous ancestry tracts, using the local recombination probabilities from the genetic map provided in ***Delaneau et al. (2019*)**. We ran a total of 19 generations, mirroring the time range from the beginning of the colonisation to present day. For computational purposes, we scaled down by a factor of 1000 the lengths and recombination rates of the genome.

For the One Pulse demographic model, we simulated a constant population size of 1000 and we set initial gene-flow proportions equivalent to the observed genetic ancestry proportions for each population ***Table 1***. For the Two Pulses model, we split the same size of migrant population from each source population in two migration waves at generation 0 and 10 as a function of the **GFR**_*gen*10_ parameter.

**Table 1.** Average proportions (%) of genetic ancestry for each population, inferred after a local ancestry analysis with RFMix.

| Ancestry | CLM | MXL | PEL | PUR | ACB | ASW |
| --- | --- | --- | --- | --- | --- | --- |
| sub-Saharan African | 8.2 | 4.3 | 3.2 | 13.8 | 88.1 | 76.5 |
| Native American | 27.7 | 50.0 | 76.1 | 15.0 | 0.5 | 4.1 |
| European | 64.1 | 45.6 | 20.7 | 71.3 | 11.4 | 19.4 |

### The continuous ancestry tract lengths profile

The continuous ancestry tract length profile is a statistic that is commonly used to date admixture events, assuming random mating. However, here we exploited the information summarised with this statistic to assess the gene-flow related to both migration and assortative mating. In addition, we included the population continuous ancestry tract length profiles of both autosomes and X chromosome to provide to the neural network information that can be used to predict sex bias. While both sexes contribute equally to the autosomal genepool, females and males contribute 2/3 and 1/3, respectively, to the X chromosome genepool. This asymmetric inheritance between autosomes and X chromosome combined with local ancestry information is highly informative of the complexity of sex-biased admixture histories ***Goldberg and Rosenberg (2015*)**.

We calculated the continuous ancestry tract length profile on simulated data, for each individual, by counting the number of tracts for each length bin, greater than or equal to the lower threshold and lower than the upper threshold, defined by a vector of break points ***b*** in a logarithmic scale, in centiMorgan (cM): 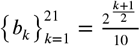. These breakpoints define a total of 22 length windows, which is a compromise of the RFMix resolution in the local ancestry analysis (0.1cM) and a limited number of windows. Then, we obtain the continuous ancestry tract length profile for each individual. Finally, we perform the mean across the individuals of the same population as the permutation-invariant function to use it as input of he neural network. For real empirical data, we run a local ancestry analysis to split the genome of each individual into the fragments inherited from Native American, European and sub-Saharan African ancestries to obtain the length of the continuous ancestrytracts (***Figure 2***A). To do this, we performed an RFMix analysis with RFMix v1.5.4 ***Maples et al. (2013)*** with the following options: -w 0.1 -G 19 -e 3. We used as target populations the six admixed populations of the Americas present in the 1000 genomes data (African Caribbeans in Barbados, ACB; African Ancestry in SW USA, ASW; Colombians in Medellín, CLM; Mexicans in Los Angeles, MXL; Peruvians in Lima, PEL; and Puerto Ricans in Puerto Rico, PUR) using the 30x coverage data ***Consortium et al. (2015); Byrska-Bishop et al. (2021)***.

To create three reference populations we first combined 1000 genomes with HGDP genomes ***Bergström et al. (2020*)**. We ran an unsupervised ***k*** = 3 ADMIXTURE analysis ***Alexander et al. (2009)***, from which we used the individuals with a proportion higher than 0.99 of one of the ancestries as reference populations for the RFMix analyses. For Native American ancestry (NAT): 6 Colombian, 12 Karitiana, 13 Maya, 13 Pima, 8 Surui, 2 MXL and 19 PEL. For European ancestry (EUR): 23 Basque, 12 Bergamoltalian, 28 French, 15 Orcadian, 28 Sardinian, 8 Tuscan, 98 CEU, 91 GBR, 98 IBS. For Sub-Saharan African ancestry (AFR): 8 BantuKenya, 8 BantuSouthAfrica, 22 Biaka, 21 Mandenka, 13 Mbuti, 6 San, 22 Yoruba, 3 ACB, 1 ASW, 99 ESN, 102 GWD, 45 LWK, 85 MSL, 107 YRI.

We considered a tract the concatenation of contiguous 0.1 cM fragments with more than 0.9 of posterior probability of being inherited from one of the three ancestries. We used the same *k*+1 break points to count the fragments in each length bin used in the simulations: 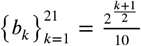 (in cM) to obtain the continuous ancestry tract length profile for each individual (***Figure 2***B). Then, we computed the mean across individuals of the same population of the continuous ancestry tract length profile to have a single matrix for each population equivalent to the output of simulations used to train the neural network. We obtain a three dimensional matrix (22×3×2) that we use as the input to the trained neural network to predict the ***AM*** and ***SB*** parameters (and ***GFR***_*gen*10_ for the Two Pulses model) (***Figure 2**C*).

### Neural network

#### Neural network architecture

We built a deep neural network comprised of four common fully-connected layers with 512, 256, 128 and 64 units, respectively, and ReLU activation functions. To avoid overfitting, we included a dropout layer with rate 0.2 after the last common layer. The network separates into five branches, each one for an independent parameter. Each branch forms a fully-connected layer with 32 units and ReLU activation functions followed by dropout with rate 0.2, and a final fully-connected output layer with a sigmoid activation function. There were 5 parameter branches for the One Pulse model (***AM***_1_, ***AM***_2_, ***AM***_2_, ***SB***_1_ and ***SB***_2_) and 3 extra parameter branches for the Two Pulses model (***GFR***_*gen*10,1_, ***GFR***_*gen*102_, ***GFR***_*geb*10,2_) (***Figure 2***C). In total the One Pulse model has 251,141 trainable weights and the Two Pulses model 263,819.

We used Adam as the optimiser and Mean Squared Error as the loss function. We rescale AM and SB from 0 to 1 to equally weight both parameters during learning. We trained the neural network for 40 epochs, a batch size of 64 with a validation split of 0.2 from the training and validation dataset. The training and validation dataset was a random 0.8 sample of the dataset comprising 10,000 matrices of the continuous ancestry tract lengths profile and we kept the remaining 0.2 for testing. We used Keras in Python to design and train the neural network ***Chollet et al. (2015)***.

All the code is available at https://github.com/massandoval/assortative_mating.

## Acknowledgments

We thank Sarah Abel and Andres Ruiz-Linares for carefully reading the manuscript and for their insightful discussion regarding the implications of the findings. We would also like to thank Flora Jay for her valuable feedback on the methods.

MF and AMS are funded are funded by The Leverhulme Research Project Grant RPG-2018-208. SM is funded in part by NIH grant R15HG011528. The content is solely the responsibility of the authors and does not necessarily represent the official views of the National Institutes of Health or other funding sources.

## Additional figures

**Figure 3–Figure supplement 1.**
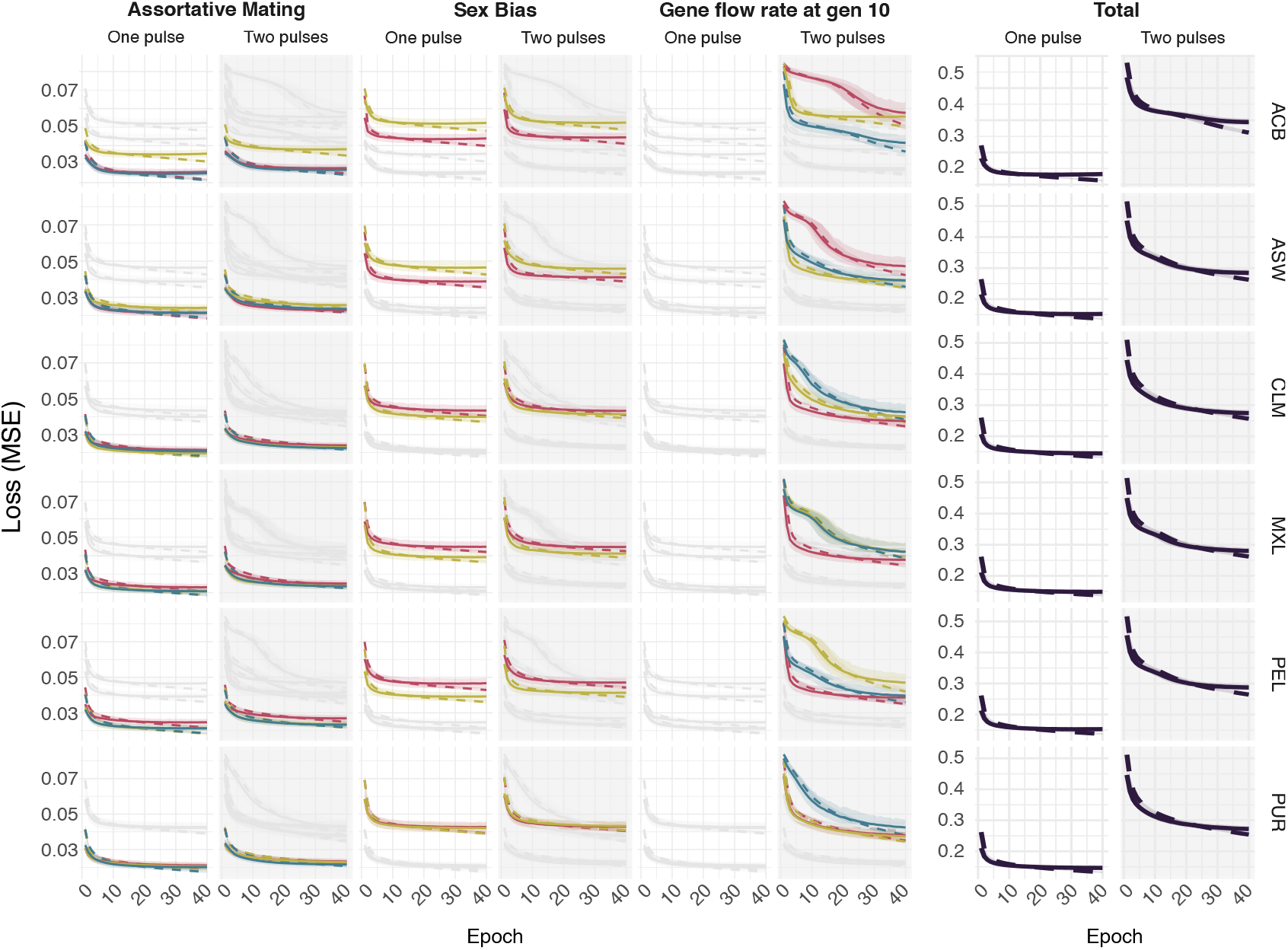
Loss function at training for the *Assortative Mating, Sex Bias, Gene flow rate at generation 10* parameters for each ancestry for both One Pulse and Two pulses models and mean values for each model. Each color is related to a different ancestry as in **Figure 3**

**Figure 3–Figure supplement 2.**
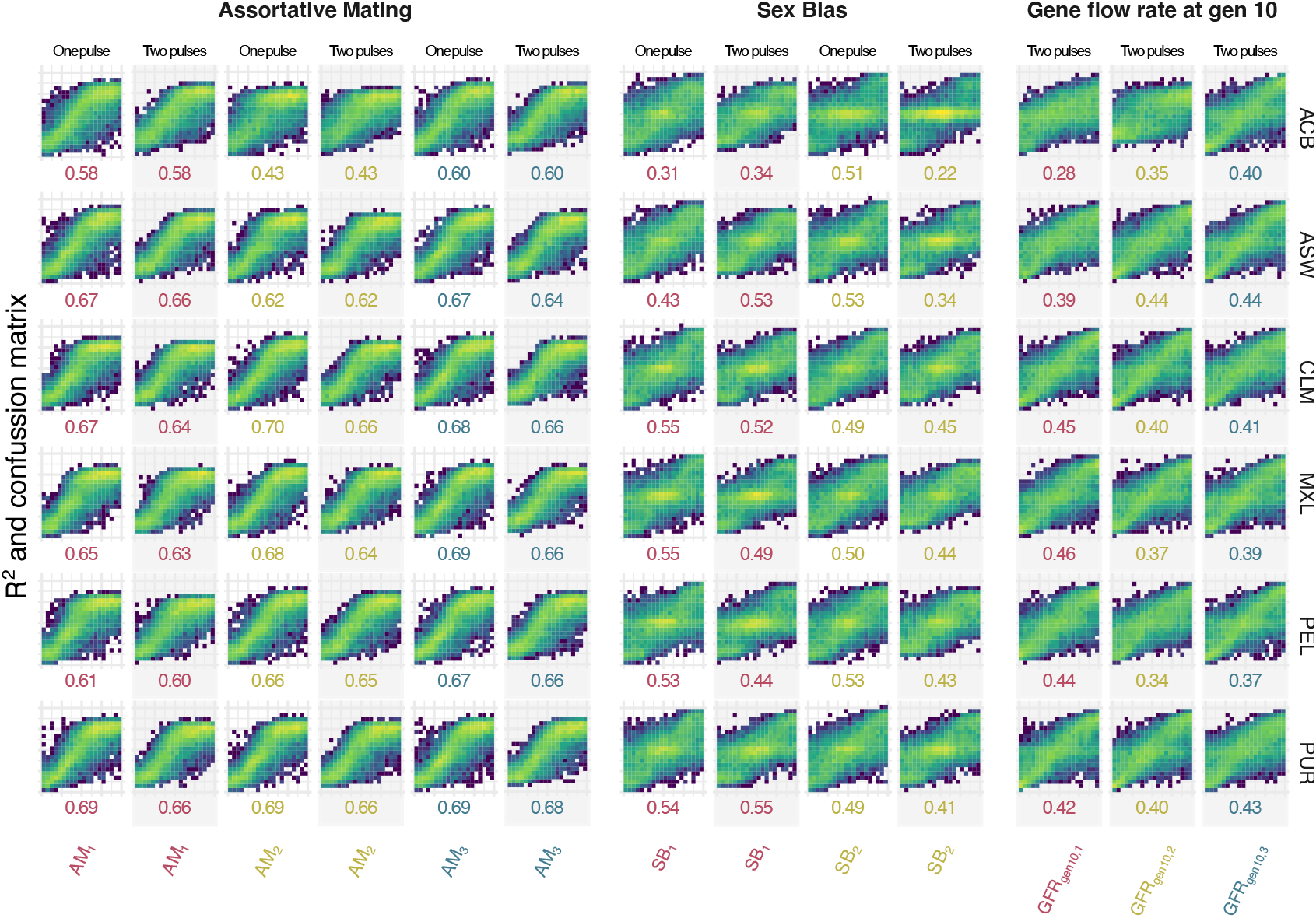
Confusion matrices and ***R***^2^ comparing true (x axis) and predicted (y axis) values at testing for *Assortative Mating, Sex Bias, Gene flow rate at generation 10* parameters for each ancestry for both One Pulse and Two pulses models. Each color is related to a different ancestry as in **Figure 3**

## Notes

### Competing Interest Statement

The authors have declared no competing interest.

